# LoRA-DR-suite: adapted embeddings predict intrinsic and soft disorder from protein sequences

**DOI:** 10.1101/2025.02.03.636253

**Authors:** Gianluca Lombardi, Beatriz Seoane, Alessandra Carbone

## Abstract

Intrinsic disorder regions (IDR) and soft disorder regions (SDR) provide crucial information on a protein structure to underpin its functioning, interaction with other molecules and assembly path. Circular dichroism experiments are used to identify intrinsic disorder residues, while SDRs are characterized using B-factors, missing residues, or a combination of both in alternative X-ray crystal structures of the same molecule. These flexible regions in proteins are particularly significant in diverse biological processes and are often implicated in pathological conditions. Accurate computational prediction of these disordered regions is thus essential for advancing protein research and understanding their functional implications. To address this challenge, LoRA-DR-suite employs a simple adapter-based architecture that utilizes protein language models embeddings as protein sequence representations, enabling the precise prediction of IDRs and SDRs directly from primary sequence data. Alongside the fast LoRA-DR-suite implementation, we release SoftDis, a unique soft disorder database constructed for approximately 500,000 PDB chains. SoftDis is designed to facilitate new research, testing, and applications on soft disorder, advancing the study of protein dynamics and interactions.

## 1 Introduction

While many proteins rely on a well-defined three-dimensional structure to carry out their functions, a substantial fraction of the proteome in any organism consists of polypeptide segments that lack a stable and reproducible, ordered structure [1, 2, 3, 4]. Remarkably, these structurally flexible or amorphous regions often retain full functionality. Such unstructured and frequently underexplored segments are known as intrinsically disordered regions (IDRs). When the intrinsic disorder spans the entirety of a protein sequence, these proteins are classified as intrinsically disordered proteins.

IDRs are typically identified using structural techniques such as circular dichroism spectroscopy. In X-ray crystallography, they appear as missing residues or highly flexible regions, often indicated by elevated B-factor values [5]. However, disorder is not always absolute—a residue may be absent in one structure yet resolved in another, reflecting a dynamic disorder-to-order transition. This inherent flexibility complicates the classification of disordered regions in protein structures. To address this challenge, the concept of *soft disorder* was introduced in [6], providing a unified framework to capture the ambiguity between ordered and disordered states. Large-scale statistical analyses of protein chains in the

Protein Data Bank (PDB) revealed a strong correlation between soft disordered regions (SDRs) and protein interaction sites [6]. These regions appear to play a crucial role in the assembly and stabilization of protein complexes [7]. Beyond this, this study provided evidence of allosteric effects, where SDRs undergo spatial rearrangements upon binding, facilitating the formation of new interaction interfaces in larger, more complex assemblies.

Given their structural flexibility, IDRs and SDRs play essential roles in molecular recognition, signaling, and regulation and are implicated in numerous diseases [8, 9, 10, 11]. Accurately identifying and characterizing these regions is crucial for advancing protein research, therapeutic development by uncovering novel biological functions and deciphering complex molecular pathways [12]. A systematic exploration of IDRs and SDRs could offer critical insights into the mechanisms of poorly understood diseases and guide therapeutic strategies targeting disordered regions and their interaction networks.

Given the fundamental role of disordered regions, ensuring the accuracy and reliability of predictive models is crucial. To this end, numerous computational tools have been developed to identify IDRs, intrinsically disordered-binding regions, and flexible segments [13, 14], with deep learning-based approaches increasingly dominating the field. To enhance their performance and practical applicability, recent efforts have systematically benchmarked these predictors using previously unresolved sequences, providing a robust assessment of their reliability and generalization capabilities [15, 16, 17]. With the rapid production of genomic and metagenomic sequences, these predictive tools serve as a crucial link between sequence data and functional annotation, delivering structural insights without the cost, time, and limitations of experimental methods.

In this work, we present LoRA-DR-suite, a novel predictor for identifying intrinsic and soft disordered regions in protein sequences. The method leverages adapted embeddings and is built on an elegantly simple deep learning architecture.

## 2 Results

### 2.1 A suite of models based on adapted embeddings

We designed (Figure 1A) a suite of classification models relying on hidden representations encoded by protein language models (PLMs). Sequences are tokenized at the single amino acid level and embedded through multiple transformer layers, generating highly-dimensional vector representations for each residue.

**Figure 1:**
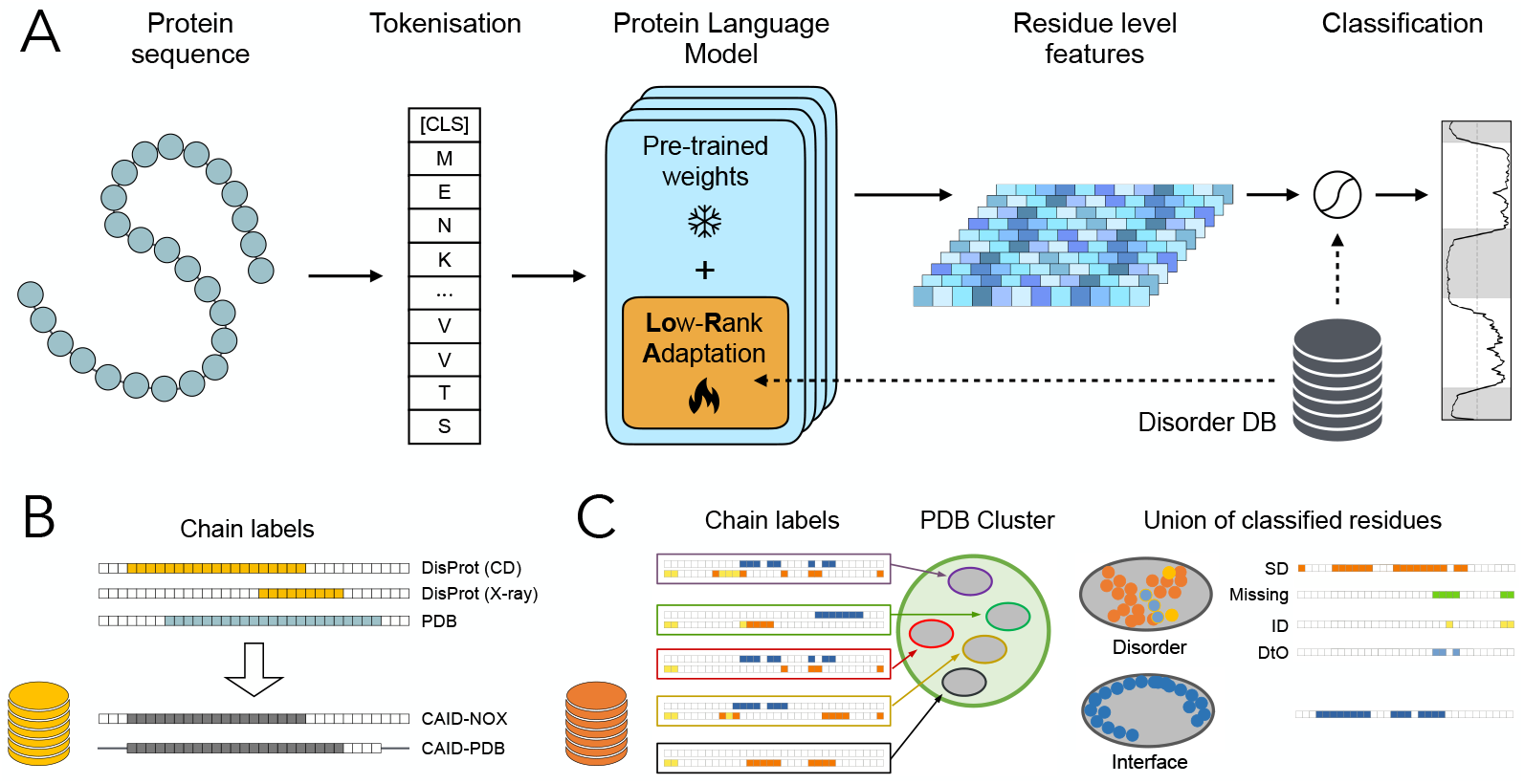
Model architecture and datasets construction. A. Taking as input a protein sequence, after tokenisation, its adapted embedding is generated by a protein language model, with frozen weights, coupled with several LoRA layers. A linear classification head with dropout is used to generate a disorder profile for the protein, where scores of disorder are computed for each residue and used to construct a disorder profile for the entire sequence. B. Three types of data are used to define intrinsic disorder. They are either coming from circular dichroism (CD) experiments, or X-ray crystallography (X-ray), or missing residues in a crystal (PDB). The CAID-PDB annotation is defined as the union of residues identified as intrinsically disordered by circular dichroism (CD) and X-ray crystallography, while the CAID-NOX annotation includes only residues identified by CD. Residues labeled as disordered or not by a method are represented by colored or white squares, respectively. Notably, missing residues in X-ray crystallographic data are unlabeled in the CAID-PDB annotation but assigned a negative label in the CAID-NOX annotation. C. In the SoftDis database, soft disordered residues (SD), missing residues (Missing), intrinsically disordered residues (ID), and disorder-to-order transition residues (DtO) are identified by clusters of chains with a very similar protein sequence (see Materials and Methods). Each protein sequence associated to a PDB structure in a cluster (left) contributes information on missing residues (yellow squares), residues with high B-factor (orange square) and interface residues (blue squares). The union of these classified residues allows to label residues as SD, Missing, ID, DtO, and interface residues for the cluster and for its sequences (right).

Exploiting the relatively large training datasets (∼ 10^5^ − 10^6^ distinct tokens for intrinsic disorder, ∼10^7^ for soft disorder), we implemented low-rank adaptation (LoRA) of the model, by introducing additional trainable layers. These layers learn low rank decomposition of large weight matrix updates [18], enabling effective modification of the model internal representations. This approach is analogous to fine-tuning but involves training only a small fraction of the model parameters. By freezing pre-trained parameters, LoRA prevents catastrophic forgetting while retaining the ability to disable adapter layers and revert to the original model. For all models, LoRA layers were added to the linear transformations in every multi-head attention module of the architecture. Details on hyperparameter choices, specific to the pre-trained model and classification task, are provided in Materials and Methods.

In this scope, the detection of disordered residues is formulated as a binary token classification task. Using pre-trained transformer architectures, we add a linear classification head with dropout, applied in parallel to the representation of each residue. As a result, LoRA-DR-suite generates a profile of intrinsic/soft disorder for the input sequence, assigning each residue a probability score indicating its likelihood of being disordered (Figure 1A, right).

In this study, we employed well-established PLMs as pre-trained models, including two versions of the Evolutionary Scale Model (ESM2) with 35M and 650M parameters [19], ProtT5 from the ProtTrans suite [20], and Ankh [21]. All models were trained using cross-entropy loss with balanced class weights to address class imbalance in the intrinsic and soft disorder datasets. These four models are integrated into the LoRA-DR-suite package.

### 2.2 Comparative Performance Analysis of LoRA-DR-suite on intrinsic disorder

The performance of LoRA-DR-suite models was evaluated against several established methods for intrinsic disorder prediction across three benchmark datasets: CAID1–DisProt [15], CAID2_NOX [16, 17], and CAID3_NOX (https://caid.idpcentral.org/challenge). These datasets were introduced by the Critical Assessment of Intrinsic protein Disorder (CAID) challenge, a biannual competition to benchmark the state-of-the-art in predicting intrinsically disordered regions in proteins: CAID1-DisProt contains the NOX subset of the first CAID round consisting of 646 protein sequences, CAID2_NOX contains 210 protein sequences from CAID2, and CAID3_NOX comprises 148 protein sequences from CAID3. Protein sequences in the NOX datasets are labeled based on information derived from circular dichroism experiments (Figure 1B). Key performance metrics include ROC AUC, F1, MCC, and PR AUC (see Materials and Methods), allowing for a comprehensive comparison of model efficacy. For each dataset, we compare with the best performing tools that participated to the corresponding CAID round.

On CAID1–DisProt test set (Table 1), ESM2_650M-LoRA achieved the best overall performance with an ROC AUC of 0.833 and a PR AUC of 0.510, indicating strong discriminative power and precision-recall trade-off. While ESM2_35M-LoRA and Ankh-LoRA followed closely in ROC AUC (0.812 and 0.819, respectively), their F1 and MCC scores showed marginal differences, underscoring the robustness of LoRA-enhanced models. Notably, these models outperformed approaches such as SPOT-Disorder2 [22] and RawMSA [23], which lagged in all metrics. This highlights the advantage of leveraging PLMs with low-rank adaptation for intrinsic disorder prediction.

**Table 1:** Results of fine-tuned models on CAID1-DisProt test set. See https://caid.idpcentral.org/challenge/results for other predictors.

| Model | ROC AUC | F1 | MCC | PR AUC |
| --- | --- | --- | --- | --- |
| ESM2_650M-LoRA | **0.833** | **0.498** | **0.394** | **0.510** |
| ESM2_35M-LoRA | 0.812 | 0.463 | 0.350 | 0.505 |
| Ankh-LoRA | 0.819 | 0.490 | 0.380 | 0.499 |
| ProtT5-LoRA | 0.817 | 0.482 | 0.373 | 0.463 |
| fIDPnn | 0.814 | 0.483 | 0.370 | 0.475 |
| fIDPlr | 0.793 | 0.452 | 0.330 | 0.421 |
| rawMSA | 0.780 | 0.445 | 0.328 | 0.413 |
| SPOT-Disorder2 | 0.760 | 0.470 | 0.349 | 0.340 |

The CAID2_NOX results (Table 2) further demonstrate the competitiveness of LoRA-enhanced PLMs. Ankh-LoRA, ProtT5-LoRA, and ESM2_650M-LoRA achieved comparable ROC AUC scores (0.837–0.838), with Ankh-LoRA slightly leading in F1 (0.548) and MCC (0.427). flDPnn2 [24] and Dispredict3 [25] performed well in ROC AUC and PR AUC, remaining competitive alternatives. Again, LoRA-enabled transformers show effectiveness in balancing precision and recall across diverse datasets.

**Table 2:** Results of fine-tuned models on CAID2_NOX test set. See also https://caid.idpcentral.org/challenge/results.

| Model | ROC AUC | F1 | MCC | PR AUC |
| --- | --- | --- | --- | --- |
| ESM2_650M-LoRA | 0.837 | 0.535 | 0.407 | 0.594 |
| ESM2_35M-LoRA | 0.813 | 0.534 | 0.406 | 0.571 |
| Ankh-LoRA | 0.838 | **0.548** | **0.427** | 0.556 |
| ProtT5-LoRA | 0.837 | 0.544 | 0.420 | 0.577 |
| fIDPnn2 | 0.838 | 0.526 | 0.414 | **0.596** |
| fIDPnn | 0.835 | 0.521 | 0.409 | 0.581 |
| Dispredict3 | **0.842** | 0.517 | 0.404 | 0.579 |
| DisoPred | 0.824 | 0.484 | 0.364 | 0.503 |
| SPOT-Disorder2 | 0.760 | 0.506 | 0.303 | 0.544 |

The CAID3_NOX benchmark (Table 3) reveals the strong performance of LoRA-enhanced models, which predict intrinsic disorder across entire protein sequences. Among them, ESM2_650M-LoRA achieved the highest scores in ROC AUC (0.880), F1 (0.649), MCC (0.524), and PR AUC (0.721), demonstrating superior predictive power. ESM2_35M-LoRA and Ankh-LoRA also performed competitively, highlighting scalability across transformer sizes. In contrast, ESMDisPred-2PDB and ESMDisPred-2, despite slightly higher PR AUC scores (0.738 and 0.732, respectively), covered only 58.9% of the total number of (positive and negative) labeled residues, focusing on selected regions rather than full-length predictions, which limits their applicability. DisoFLAG-IDR and flDPnn3a showed lower ROC AUC and PR AUC scores (0.857 and 0.851, respectively) despite their full-length coverage, further underscoring the advantage of LoRA-adapted PLM-based strategies.

**Table 3:** Results of fine-tuned models on the CAID3_NOX test set. F1 and MCC scores were calculated for comparative tools as they were not provided on the official results page https://caid.idpcentral.org/challenge/results. Visit the same address for comparisons with additional predictors.

| Model | ROC AUC | F1 | MCC | PR AUC | Cov % |
| --- | --- | --- | --- | --- | --- |
| ESM2_650M-LoRA | **0.880** | **0.649** | **0.523** | 0.721 | 100 |
| ESM2_35M-LoRA | 0.868 | 0.645 | 0.518 | 0.689 | 100 |
| Ankh-LoRA | 0.861 | 0.637 | 0.507 | 0.684 | 100 |
| ProtT5-LoRA | 0.862 | 0.628 | 0.494 | 0.698 | 100 |
| ESMDisPred-2PDB | 0.868 | 0.572 | 0.467 | **0.738** | 58.9 |
| ESMDisPred-2 | 0.859 | 0.603 | 0.480 | 0.732 | 58.9 |
| rawMSA-disorder | 0.858 | 0.504 | 0.399 | 0.640 | 100 |
| DisoFLAG-IDR | 0.857 | 0.629 | 0.499 | 0.694 | 84.5 |
| fIDPnn3a | 0.851 | 0.587 | 0.436 | 0.687 | 100 |

Across all benchmarks, ESM2_650M-LoRA consistently demonstrated strong performance, particularly on the dataset CAID3_NOX, where additional data from CAID1 and CAID2 were used for training (see Materials and Methods). The addition of LoRA layers significantly enhances the capacity of pre-trained language models to predict intrinsically disordered residues with precision and robustness, even under class imbalances. Models like SPOT-Disorder2 and RawMSA lagged behind, while hybrid models such as flDPnn showed competitive but narrower advantages.

### 2.3 Extending the definition of disorder through the database SoftDis for soft disorder

The concept of soft disorder was introduced in [6] as a general term for regions in a protein identified as flexible (characterized by a high B-factor) or intermittently missing across different X-ray crystal structures of the same sequence. Formally, this definition is derived from an extensive analysis of clusters of alternative structures for the same protein sequence in the Protein Data Bank (PDB): based on the same procedure introduced in [6], we newly analyzed 229,370 crystallographic structures and clustered their 484,044 chains into 64,285 clusters to create the SoftDis database. A residue is classified as soft-disordered if it is flexible or missing in at least one crystal structure within a cluster, excluding residues that are consistently missing across all structures. Based on this definition, we statistically demonstrated a significant correlation between SDRs and interface sites observed in protein complexes of identical sequences [6]. This analysis also revealed that disordered regions often undergo structural shifts upon binding, facilitating the formation of new interfaces that arise during the assembly pathway [7]. The SoftDis database is available on the HuggingFace datasets Hub (see Data and software availability). It includes labels for representative sequences in clusters, covering soft disorder, missing residues, and interfaces as identified in the PDB. Reduced datasets based on similarity clustering and non-redundant splits designed for learning tasks are also included, along with detailed data for chains within each cluster.

### 2.4 Soft disorder classification

Results on intrinsic disorder classification demonstrate the ability of PLMs fine-tuning in the identification of disorder markers, that we wanted to extend to the broader definition of soft disorder. In this scope, we evaluated the same suite of models on the new soft disorder dataset that we constructed, whose specific details are discussed in the Materials and Methods Section. For learning and evaluation, we first reduced redundancy of training data by clustering representative sequences for each PDB cluster at 50% sequence identity, then randomly split remaining sequences into train, validation and test sets. Sequences in validation and test sets were further filtered at a 30% sequence identity threshold, to ensure higher generalizability of evaluation performances. For each fine-tuned PLM of LoRA-DR-suite, the hyperparameters search was performed according to the same settings as in the case of intrinsic disorder, but using a reduced sample (15%) of all training data for feasibility. Models with selected parameters were then re-trained on the full dataset.

Table 4 provides a performance comparison of LoRA-DR-suite models for predicting soft disordered residues.

**Table 4:** Results of LoRA-DR-suite models trained on soft disorder on the SoftDis test set.

| Model | ROC AUC | F1 | MCC | PR AUC |
| --- | --- | --- | --- | --- |
| ESM2_650M-LoRA | **0.839** | **0.655** | **0.476** | **0.729** |
| ESM2_35M-LoRA | 0.812 | 0.626 | 0.430 | 0.690 |
| Ankh-LoRA | 0.831 | 0.646 | 0.468 | 0.716 |
| ProtT5-LoRA | 0.823 | 0.638 | 0.451 | 0.702 |

The results highlight a trade-off between model complexity and predictive performance: ESM2_650M-LoRA, with its larger size and higher computational demand, achieves the best results, while smaller models like ESM2_35M-LoRA deliver slightly lower performance but remain computationally efficient. Ankh-LoRA offers a balanced middle ground between these extremes. These results parallel those obtained for intrinsic disorder classification, underscoring the predictive power of adapted embeddings in recognizing disorder patterns. Notably ESM-derived models demonstrate higher success rates despite their lower architectural complexity.

To illustrate the performance of LoRA-DR-suite on soft disorder predictions, we analysed the Bacillus subtilis Oxalate Decarboxylase OxdC protein (PDB ID 2UYA). Experimental data from the SoftDis database were used to compare the union of interface residues (Figure 2A) and the union of soft disordered residues (Figure 2B) computed on all PDB complexes containing this chain, to the predicted soft disorder residues (Figure 2C). Notably, the predicted soft disordered residues show a high correlation with the experimental data. Additionally, low pLDDT values from the AlphaFold structural model of the protein (Figure 2D) align with the predicted soft disordered residues [26], further supporting the robustness of the LoRA-DR-suite’s predictions.

**Figure 2:**
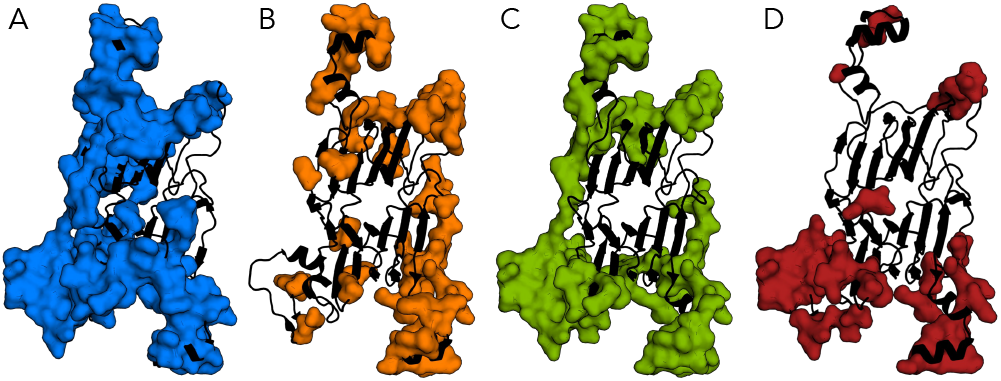
Structural analysis of the Bacillus subtilis Oxalate Decarboxylase OxdC protein. (A) Residues involved in binding sites across any complex in the SoftDis cluster containing PDB ID 2UYA are highlighted in dark blue. (B) Soft disordered residues identified in all PDB complexes, as in A, are shown in orange. (C) Residues predicted as soft disordered by ESM2_650M-LoRA are highlighted in green. (D) Residues with pLDDT scores below the chain’s average are highlighted in red.

### 2.5 Linking soft and intrinsic disorder with pLDDT

We conducted a large scale analysis on the 2009 chains from our SoftDis testing dataset that have structural models available in the AlphaFold database. This analysis reveals that lower pLDDT values—indicating lower structural confidence or higher disorder—correlate positively with soft disorder predictions and the frequency of soft disorder observed in similar protein chains (Figure 3A). These results suggest that regions with low pLDDT scores align well with both predicted and observed soft disorder patterns, effectively capturing structural variability and flexibility. Figure 3B further shows that pLDDT scores align with LoRA-DR-suite predictions for both intrinsic and soft disorder. However, the correlation is stronger for soft disorder, highlighting key distinctions in how these two disorder types relate to structural confidence [27].

**Figure 3:**
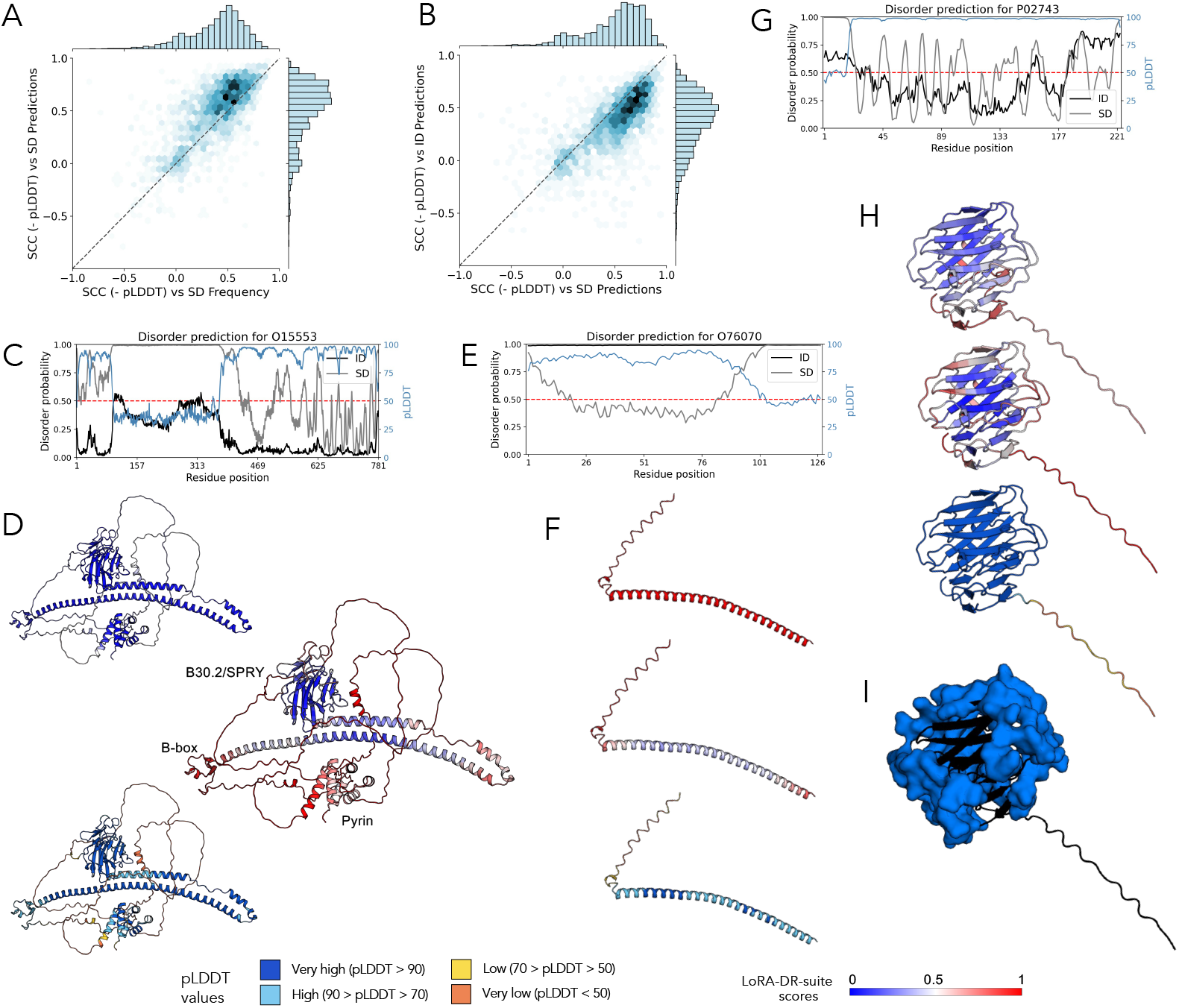
Structural analysis of the MEFV, Synuclein and SAMP human proteins. (A) Heatmap of similarity (measured by the Spearman Correlation Coefficient) between pLDDT scores and ESM2_650M-LoRA scores for soft disorder predictions, and between pLDDT scores and the frequency of a residue being soft disordered within the PDB cluster in the SoftDis database. The correlations are calculated on the negative of the pLDDT scores. (B) As in A, heatmap between pLDDT scores and ESM2_650M-LoRA scores of intrinsic disorder, and pLDDT and ESM2_650M-LoRA scores of soft disorder predictions. (C) Profiles of ESM2_650M-LoRA scores of soft disorder, ESM2_650M-LoRA scores of intrinsic disorder, and pLDDT for the MEFV protein (UniProt ID O15553). (D) MEFV structural model in AlphaFold Database colored with ESM2_650M-LoRA intrinsic disorder scores (top), ESM2_650M-LoRA soft disorder scores (middle), and pLDDT confidence scores (bottom). (E) Profiles of ESM2_650M-LoRA scores of soft disorder, ESM2_650M-LoRA scores of intrinsic disorder, and pLDDT for the Synuclein protein (UniProt ID O76070). (F) Synucleins structural model in AlphaFold Database colored with ESM2_650M-LoRA intrinsic disorder scores (top), ESM2_650M-LoRA soft disorder scores (middle), and pLDDT confidence scores (bottom). (G) Profiles of ESM2_650M-LoRA scores of soft disorder, ESM2_650M-LoRA scores of intrinsic disorder, and pLDDT for the Serum amyloid P-component protein (SAMP; UniProt ID P02743). (H) AlphaFold model structure of the SAMP protein with residues colored by predicted scores of intrinsic disorder (top), predicted scores of soft disorder (middle), and pLDDT values (bottom). Corresponding color bars are shown on the right. (I) PDB structure 1GYK, chain A, with residues highlighted in blue to indicate interface residues identified in SoftDis. These residues were computed using the cluster represented by chain C of PDB structure 1SAC, which contains 90 chains.

Figures 3C and 3D focus on the MEFV (Mediterranean Fever) human protein gene, an important modulator of innate immunity. The soft disorder and pLDDT profiles exhibit a strong Spearman correlation (0.865 with -pLDDT), with soft disorder effectively capturing flexibility differences across annotated domains that are not detected as intrinsic disorder. For example, the pyrin domain (1–92) and B-box domain (370–412) are identified as highly flexible, whereas the B30.2/SPRY domain (580–775) appears much more stable. Additionally, the long helices (420–582), required for homotrimerization and pyroptosome induction, exhibit variable flexibility: stable at the center (low soft disorder, high pLDDT) but highly flexible at the helix extremes (high soft disorder, low pLDDT). These patterns are undetected by intrinsic disorder predictions, which show a weaker correlation with -pLDDT (0.796).

Figures 3E and 3F analyze the Synuclein protein. While our intrinsic disorder predictor classifies it as highly disordered, the soft disorder profile reveals a more detailed perspective: the helix’s extremes are identified as highly flexible, while the central region is stable. -pLDDT aligns strongly with soft disorder predictions (correlation of 0.816), while its correlation with intrinsic disorder predictions is weaker (0.505), underscoring the differences between these two disorder types.

Figures 3G and 3H analyze the Serum amyloid P-component protein. This protein exhibits consistently high pLDDT values across its sequence, with correlations of 0.591 and 0.829 observed between -pLDDT and intrinsic disorder and soft disorder, respectively. The soft disorder profile identifies several regions with high scores, indicative of flexible zones. These regions are localized on the protein surface (Figure 3H, center) and correspond to interaction sites, as highlighted by the blue residues in Figure 3I, derived from the SoftDis database.

These observations underscore the relevance of soft disorder predictions in identifying structurally flexible, functionally important regions, involved in protein-protein interactions. The strong correlation with -pLDDT further highlights the potential of combining structural confidence scores with soft disorder profiles to enhance our understanding of protein flexibility, structural variability and interaction dynamics.

### 2.6 Adapted embeddings influence on attention scores

LoRA-DR-suite models exploit adapted embeddings to extract and distinguish fundamental differences between disorder signals. To explore whether intrinsic and soft disorder residues are predominantly influenced by global or local effects learned through the adapter layers, we utilized contact prediction as an indirect method to analyze variations in attention scores caused by model fine-tuning [28].

We performed a large scale analysis of variations in predicted contact maps across 2,512 chains from the SoftDis test set, excluding those longer than 1,024 residues, to assess the impact of LoRA layers compared to the base model ESM2_650M. Adapted embeddings were generated by training with information from either intrinsic or soft disorder, while contact probabilities were derived from the pre-trained ESM2_650M contact prediction head on attention outputs. Figure 4A shows that ESM2_650M-LoRA for intrinsic disorder induces variations primarily at very short ranges. In contrast, training on soft disorder leads to variations observed at much greater distances, extending up to a third of the protein’s length. Although the number of long-range variations decreases with increasing distance, they remain consistently present. Figure 4B presents an in-depth analysis of the phosphate system positive regulator protein PHO4 (chain B), whose dimer is known to form a complex with DNA. The variability in contact probabilities is significantly more pronounced when training on soft disorder, with notable differences observable (see blue squares) at short range compared to embeddings trained on intrinsic disorder. For intrinsic disorder, adapted embeddings tend to loose contact information at larger ranges (see red squares), a phenomenon also present for soft disorder but to a lesser extent. In Figure 4C, positions with enhanced contact scores for the model trained on soft disorder are shown. High scores near the principal diagonal (blue) are mostly found in coiled regions, while those at larger distances reveal contacts with a partner chain in the dimeric complex. Notably, residues predicted to contact alanine at position 20 (pink) and leucine at position 27 (magenta) are highlighted. These results emphasize the unique influence of soft disorder training on capturing contact probabilities through adapted embeddings.

**Figure 4:**
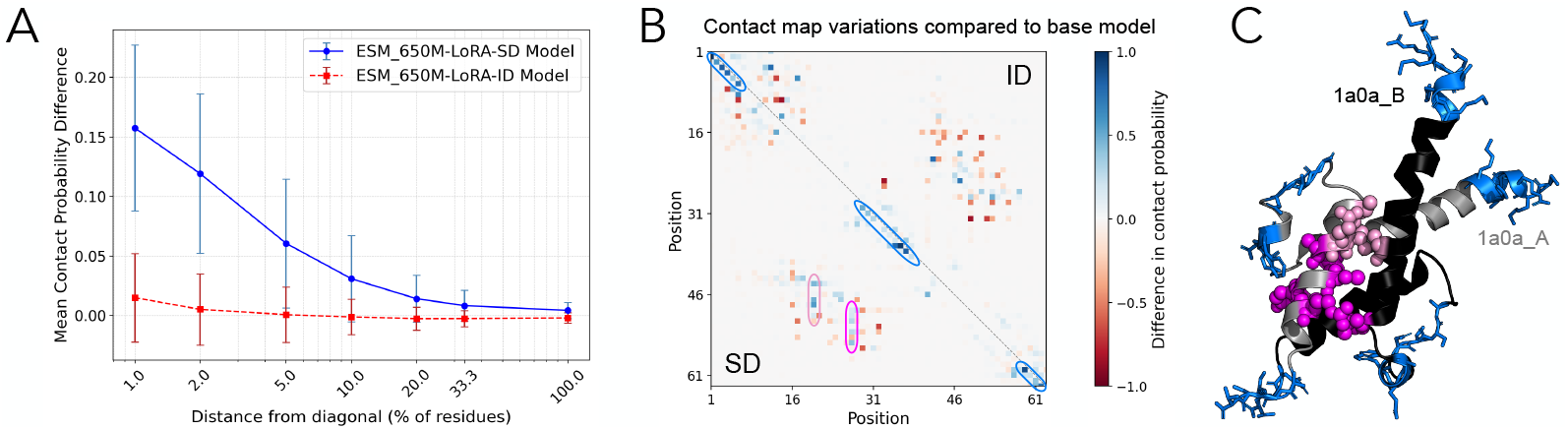
Variations in predicted contacts maps induced by LoRA layers. (A) Average variation in contacts probability scores returned by ESM2_650M contact prediction head, between models fine-tuned on soft disorder (ESM2_650M-LoRA-SD) and intrinsic disorder (ESM2_650M-LoRA-ID) with respect to the ESM2_650M base model, at different distances from the principal diagonal (measured as fraction of total protein length). (B) Example of contact map variations induced by fine-tuned models on soft disorder (SD, below diagonal) and intrinsic disorder (ID, above diagonal) for PDB chain 1A0A_B. (C) Enhanced predicted contacts from soft disorder fine-tuning are plotted on the dimeric PDB structure (1A0A; DNA chains are removed). Positions with increased scores near the diagonal are colored in blue on both chains. Residues showing enhanced contacts with positions 20 and 27 from chain 1A0A_B are colored in pink and magenta, respectively, in chain 1A0A_A.

## 3 Materials and Methods

### 3.1 Intrinsic disorder datasets

For model selection and training, we used the dataset of [29], extracted from the DisProt 7.0 database [30, 31, 32] following the same preprocessing procedure. The dataset originally comprised 745 experimentally labeled sequences, which were divided into training (445 sequences), validation (100 sequences) and test (200 sequences) sets. To ensure low sequence similarity between training and test data, the combined training and test sets were clustered at 25% sequence identity using the CD-HIT algorithm [33]. Sequences from the test set that belonged to the same clusters as training sequences were removed, resulting in a final test set of 176 sequences with low sequence identity relative to the training set.

To evaluate the generalization capabilities of the models, we also used datasets from the first, second and third editions of the CAID challenge. The CAID1-DisProt dataset contains 646 protein sequences that were unknown to the contestants before the evaluation, and whose labels follow the DisProt format. Proteins were sourced from diverse species, including humans and other mammals (368 sequences) as well as prokaryotes (77 sequences). Similarly, the CAID2_NOX dataset consists of 210 sequences from the Disorder_NOX subset of the second challenge edition, while the CAID3_NOX includes 148 sequences from the Disorder_NOX subset of the third edition. Moreover, in the CAID3 evaluation, we added all sequences from CAID1-DisProt and CAID2_NOX, as well as the initially held out set, to the training data, while keeping the original 100 sequences for validation. Sequences in the training and validation sets were truncated to a maximum length of 1024 residues in all the experiments. See the legend of Figure 1B for the definition of intrinsic disorder, CAID-NOX and CAID-PDB.

### 3.2 Construction of the soft disorder database SoftDis

To construct the SoftDis database, we retrieved protein structures from the PDB archive (snapshot as of 15-10-2024) and clustered sequences using MMSeqs2 [34, 35] at 90% sequence identity and 90% coverage. This process yielded 64,285 clusters encompassing a total of 484,044 chains belonging to 229,376 structures. The average number of structures per cluster was 7.53, with a median of 3. Nearly half of the clusters (31,412) contained only 1 or 2 sequences, while the largest cluster included 1,727 homologs.

The representative chain for each cluster was selected from the experimental structure with the highest resolution or best R-value. For protein complexes, individual chains were analyzed separately and assigned to their respective clusters.

Residues in each chain were labeled as missing if annotated under REMARK 465 in the PDB file. A residue was classified as soft disordered if it was either missing or had a normalized B-factor 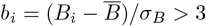, where *B*_*i*_ represents the B-factor of the *C*_*α*_ atom, and 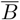 and *σ*_*B*_ are the mean and standard deviation of *B*_*i*_ values within the chain. Additionally, residues were labeled as interface if they participated in protein-protein or protein-DNA/RNA interactions. Protein-protein binding sites were identified using the INTerface Builder tool [36], where residue contacts are defined by *C*_*α*_ atoms within 5 Å. Protein-DNA/RNA binding sites were determined as residues showing decreased accessible surface area, measured using Naccess [37] with a 1.4 Å probe, upon binding.

For each site in the representative sequence of a cluster, we recorded the number of times it was labeled as soft disordered across all chains in the cluster, excluding sites consistently labeled as missing. Similarly, we noted the number of times each site was labeled as interface. Figure 1C illustrates the process used to assign labels for soft disordered residues, missing residues, intrinsically disordered residues, and disorder-to-order transition residues in the SoftDis database.

To adapt the dataset for machine learning pipelines, we removed sequences shorter than 20 amino acids and longer than 2,048, and reduced redundancy by further clustering representative sequences at 50% sequence identity and 80% coverage. The resulting 38,218 sequences were then randomly split into train, validation and test set, with ratios of 0.7, 0.1 and 0.2 respectively. Validation and test sets were then pruned at different levels of sequence identity with respect to the train set, at 0.9, 0.7, 0.5 and 0.3 thresholds, allowing for increasingly more difficult evaluation scenarios. For training and evaluation, we used the version of the dataset with test set pruning at 0.3 threshold, comprising 26,752 sequences for training, 1155 for validation and 2523 for testing.

To generate binary labels for soft disorder classification, residues are assigned a value of 1 if they appear as soft disordered in at least one sequence in the cluster, and 0 otherwise. The average percentage of soft-disordered residues per sequence is approximately 32% in all data splits. Additionally, we provide the fraction of chains where each residue is identified as soft disordered, enabling model uncertainty quantification. A similar analysis applied to residues at protein-protein or protein-DNA/RNA interaction sites supports multi-label classification for each position.

### 3.3 Evaluation metrics

To properly evaluate LoRA-DR-suite performance we rely on a ground truth set by reference datasets, depending on the experiment, and the following quantities: known disordered residues (intrinsic or soft) that are identified by LoRA-DR-suite (true positives, TP), disordered residues identified by LoRA-DR-suite which are not known (false positives, FP), disordered residues that are not found by LoRA-DR-suite but are known (false negatives, FN), and disordered residues that are not known and are not detected by LoRA-DR-suite (true negatives, TN).

Due to the label imbalance, we considered the following four standard metrics of performance:

– Area under the Receiver Operating Characteristic curve (ROC AUC), that shows the trade-off between true positive rate (TPR) and false positive rate (FPR) across different decision thresholds.
– Matthews Correlation Coefficient:
– MCC = (*TP* · *TN* − *FP* · *FN*) */K*, where 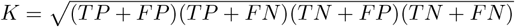
– F1 score = 2 *TP/*(2 *TP* + *FP* + *FN*)
– Area under the Precision Recall curve (PR AUC), computed as average precision, where the precision-recall metric shows the tradeoff between precision and recall for different thresholds, with recall (sensitivity) = *TP/*(*TP* + *FN*) and precision (positive predictive value) = *TP/*(*TP* + *FP*).

### 3.4 PLMs fine-tuning and hyperparameters selection

Pre-trained PLMs checkpoints used for our evaluation were taken from the HuggingFace Hub, that allows easy customization for parameters efficient fine-tuning strategies, such as Low Rank Adaptation (LoRA) of attention layers. Hyperparameters optimization was run with HuggingFace Trainer class and Optuna backend [38], with the following possible choices in all experiments:

– Discrete parameters:
  – LoRA rank *r* ∈ *{*8, 16, 32*}*;
  – dropout probability *p* ∈ *{*0.1, 0.2, 0.3*}*;
  – scheduler warm up ratio ∈ *{*0.1, 0.2*}*;
  – optimizer weight decay ∈ *{*0.1, 0.01, 0.001*}*.
– Maximum learning rate *lr* ∈ [10^−5^, 10^−3^], with uniform probability on logarithmic scale.
– Attention layers with LoRA adapters, either {*Q, K, V*} or {*Q, V*}.

Optimizer was set to AdamW [39] with default parameters and cosine scheduler decay. We used an effective batch size of 8 for training on Disprot 7.0, and of 16 for training on the full intrinsic disorder and soft disorder datasets.

For each experiment, the models with the highest ROC AUC on the validation set were selected from 40 trials using random parameter search (Table 5). Models were trained for up to 10 epochs, with early stopping regularization based on validation ROC AUC and patience set to 2. Training and evaluations were performed on a single computing node equipped with an Intel Xeon Gold 6330 processor (4 cores, 64 GB RAM reserved) and a NVIDIA A100 PCIe GPU with 80GB RAM.

**Table 5:** Hyperparameters selected for each model, and ROC AUC score obtained on the relative validation set. DisProt7 suffix refers to models evaluated on the first and second edition of CAID, thus trained only on the DisProt 7.0 subset, while models with ID (SD) suffix are trained on the full intrinsic disorder (soft disorder) datasets.

| Model | LoRA $r$ | Dropout $p$ | LoRA layers | Learning rate | Warm up ratio | Weight decay | ROC AUC |
| --- | --- | --- | --- | --- | --- | --- | --- |
| ESM2_650M-LoRA-DisProt7 | 16 | 0.1 | $Q, V$ | $4.11 \cdot 10^{-4}$ | 0.1 | 0.1 | 0.842 |
| ESM2_35M-LoRA-DisProt7 | 32 | 0.3 | $Q, V$ | $7.09 \cdot 10^{-4}$ | 0.1 | 0.1 | 0.836 |
| Ankh-LoRA-DisProt7 | 16 | 0.3 | $Q, V$ | $5.64 \cdot 10^{-4}$ | 0.1 | 0.001 | 0.831 |
| ProtT5-LoRA-DisProt7 | 16 | 0.3 | $Q, V$ | $9.74 \cdot 10^{-4}$ | 0.1 | 0.01 | 0.853 |
| ESM2_650M-LoRA-ID | 32 | 0.1 | $Q, V$ | $2.08 \cdot 10^{-4}$ | 0.1 | 0.001 | 0.858 |
| ESM2_35M-LoRA-ID | 16 | 0.3 | $Q, K, V$ | $9.37 \cdot 10^{-4}$ | 0.2 | 0.001 | 0.848 |
| Ankh-LoRA-ID | 32 | 0.3 | $Q, V$ | $2.41 \cdot 10^{-4}$ | 0.2 | 0.01 | 0.836 |
| ProtT5-LoRA-ID | 8 | 0.2 | $Q, V$ | $7.73 \cdot 10^{-4}$ | 0.1 | 0.1 | 0.847 |
| ESM2_650M-LoRA-SD | 32 | 0.3 | $Q, V$ | $5.49 \cdot 10^{-4}$ | 0.2 | 0.01 | 0.844 |
| ESM2_35M-LoRA-SD | 16 | 0.1 | $Q, V$ | $5.80 \cdot 10^{-4}$ | 0.1 | 0.01 | 0.815 |
| Ankh-LoRA-SD | 8 | 0.1 | $Q, K, V$ | $8.09 \cdot 10^{-4}$ | 0.1 | 0.01 | 0.837 |
| ProtT5-LoRA-SD | 16 | 0.2 | $Q, V$ | $9.93 \cdot 10^{-4}$ | 0.1 | 0.001 | 0.826 |

## 4 Discussion

By incorporating LoRA adaptations into PLMs, the LoRA-DR-suite achieves performance that matches or surpasses state-of-the-art methods, positioning itself as a highly competitive tool for intrinsic disorder prediction. Its versatility allows it to effectively address both intrinsic and soft disorder, broadening its applicability and paving the way for new research directions in the field.

### Why a Predictor of Soft Disorder?

By predicting regions of flexibility or transient disorder across ensembles of related proteins, LoRA-DR-suite models provide valuable insights in the following areas:

#### 1. Identification of protein interactions sites

Protein interfaces are critical regions, often sensitive to mutations and linked to pathogenic variants. IDRs enable diverse binding modes, facilitating regulation, recognition, and signaling through high-specificity, low-affinity interactions [8]. SDRs further enrich interface complexity, with residues of defined interface roles—transient or permanent—that can dynamically shift between structured and disordered states [7]. Hence, differentiating intrinsic disorder, soft disorder, and other residues is key to understanding protein interfaces, and the integration of these concepts into models of protein-protein interactions and protein dynamics holds the potential for significant breakthroughs.

#### 2. Link between disorder and assembly pathways

Soft disorder reveals how structural flexibility shapes dynamic protein interactions and functional networks [7]. To this end, soft disorder-informed predictions can serve as a foundation for designing network-targeted therapeutics that leverage the unique properties of disordered regions. In addition, the connection between soft disorder and assembly could reveal how proteins exploit flexibility and transient disorder for multivalent interactions.

#### 3. Guidance for experimental studies

Accurate and fast sequence-based predictions help to efficiently screen large sequence datasets and prioritize candidate residues for validation, reducing reliance on costly and time-intensive methods for structural determination.

### Expanding protein disorder research through multiple disorders

Integrating intrinsic disorder—defined via circular dichroism on isolated protein chains—with soft disorder—identified through flexibility and missing residues across similar X-ray structures—broadens disorder research in many ways.

Indeed, the determination of factors causing the overlapping and divergence between the two notions of disorder, favored by LoRA-DR-suite predictions and SoftDis database, could reveal their link to specific functions like signaling, regulation, and scaffolding, indicating distinct roles of intrinsic and soft disorder in biological processes, and uncover the interplay between structural and environmental factors that shape disorder dynamics. Furthermore, exploring the evolutionary constraints that influence intrinsic and soft disorder and their possible variations across species or protein families may reveal distinct selective pressure shaping protein functional diversity [40].

Future research in these directions is critical for our understanding of how intrinsic and soft disorder contribute to human diseases, where disruptions in disorder-mediated interactions drive pathological states [8]. Studying intrinsically-disordered and hybrid proteins—comprising ordered domains and functional IDRs with residues that are missing, disordered-to-ordered, consistently flexible, or flexible-to-rigid—could provide key insights into disease mechanisms and novel therapeutic targets.

### Repurposable adapted embeddings

While not explored in this work, the adapted embeddings produced by the models can be extracted, analyzed, or repurposed for other tasks related to intrinsic or soft disorder in proteins. The prediction of conditions or partners that induce disorder-to-order transitions in soft disorder regions is a promising research direction that would likely benefit from LoRA-DR-suite models, connecting predictive modeling with functional insights.

## 5 Data and software availability

LoRA-DR-suite models and SoftDis dataset are available at https://huggingface.co/CQSB, with sample scripts for model deployment, dataset loading and processing. Scripts for hyperparameters optimization and models testing are available at http://gitlab.lcqb.upmc.fr/lombardi/LoRA-DR-suite.

## 6 Aknowledgements

Institut Universitaire de France (AC); Sorbonne Center for Artificial Intelligence (GL). This work was performed using HPC resources from Sorbonne University GPU clusters at the LIP6 laboratory (UMR 7606, Sorbonne University-CNRS).

We thank Alessio Corrado, Damien Legros and Maya Czeneszew for their participation in the initial steps of the project.

## References

[1] Peter Tompa. Intrinsically unstructured proteins. Trends in biochemical sciences, 27(10):527–533, 2002.

[2] A Keith Dunker, Pedro Romero, Zoran Obradovic, Ethan C Garner, and Celeste J Brown. Intrinsic protein disorder in complete genomes. Genome informatics, 11:161–171, 2000.

[3] Bálint Mészáros, István Simon, and Zsuzsanna Dosztányi. Prediction of protein binding regions in disordered proteins. PLoS computational biology, 5(5):e1000376, 2009.

[4] Gang Hu, Kui Wang, Jiangning Song, Vladimir N Uversky, and Lukasz Kurgan. Taxonomic landscape of the dark proteomes: Whole-proteome scale interplay between structural darkness, intrinsic disorder, and crystallization propensity. Proteomics, 18(21-22):1800243, 2018.

[5] Zhoutong Sun, Qian Liu, Ge Qu, Yan Feng, and Manfred T Reetz. Utility of B-factors in protein science: interpreting rigidity, flexibility, and internal motion and engineering thermostability. Chemical reviews, 119(3):1626–1665, 2019.

[6] Beatriz Seoane and Alessandra Carbone. The complexity of protein interactions unravelled from structural disorder. PLOS Computational Biology, 17(1):e1008546, 2021.

[7] Beatriz Seoane and Alessandra Carbone. Soft disorder modulates the assembly path of protein complexes. PLOS Computational Biology, 18(11):e1010713, 2022.

[8] Vladimir N Uversky. Intrinsic disorder, protein–protein interactions, and disease. Advances in protein chemistry and structural biology, 110:85–121, 2018.

[9] Ana M Melo, Juliana Coraor, Garrett Alpha-Cobb, Shana Elbaum-Garfinkle, Abhinav Nath, and Elizabeth Rhoades. A functional role for intrinsic disorder in the tau-tubulin complex. Proceedings of the National Academy of Sciences, 113(50):14336–14341, 2016.

[10] Kumlesh K Dev, Katja Hofele, Samuel Barbieri, Vladimir L Buchman, and Herman van der Putten. Part II: α-synuclein and its molecular pathophysiological role in neurodegenerative disease. Neuropharmacology, 45(1):14–44, 2003.

[11] Lilia M Iakoucheva, Celeste J Brown, J David Lawson, Zoran Obradović, and A Keith Dunker. Intrinsic disorder in cell-signaling and cancer-associated proteins. Journal of molecular biology, 323(3):573–584, 2002.

[12] A Keith Dunker and Vladimir N Uversky. Drugs for ‘protein clouds’: targeting intrinsically disordered transcription factors. Current opinion in pharmacology, 10(6):782–788, 2010.

[13] Fanchi Meng, Vladimir N Uversky, and Lukasz Kurgan. Comprehensive review of methods for prediction of intrinsic disorder and its molecular functions. Cellular and Molecular Life Sciences, 74:3069–3090, 2017.

[14] Lukasz Kurgan, Gang Hu, Kui Wang, Sina Ghadermarzi, Bi Zhao, Nawar Malhis, Gábor Erdo?s, Jörg Gsponer, Vladimir N Uversky, and Zsuzsanna Dosztányi. Tutorial: a guide for the selection of fast and accurate computational tools for the prediction of intrinsic disorder in proteins. Nature Protocols, 18(11):3157–3172, 2023.

[15] Marco Necci, Damiano Piovesan, and Silvio CE Tosatto. Critical assessment of protein intrinsic disorder prediction. Nature methods, 18(5):472–481, 2021.

[16] Alessio Del Conte, Mahta Mehdiabadi, Adel Bouhraoua, Alexander Miguel Monzon, Silvio CE Tosatto, and Damiano Piovesan. Critical assessment of protein intrinsic disorder prediction (CAID)-Results of round 2. Proteins: Structure, Function, and Bioinformatics, 91(12):1925–1934, 2023.

[17] Alessio Del Conte, Adel Bouhraoua, Mahta Mehdiabadi, Damiano Clementel, Alexander Miguel Monzon, Silvio CE Tosatto, and Damiano Piovesan. CAID prediction portal: a comprehensive service for predicting intrinsic disorder and binding regions in proteins. Nucleic Acids Research, 51(W1):W62–W69, 2023.

[18] Yu Yu, Chao-Han Huck Yang, Jari Kolehmainen, Prashanth G Shivakumar, Yile Gu, Sungho Ryu Roger Ren, Qi Luo, Aditya Gourav, I-Fan Chen, Yi-Chieh Liu, et al. Low-rank adaptation of large language model rescoring for parameter-efficient speech recognition. In 2023 IEEE Automatic Speech Recognition and Understanding Workshop (ASRU), pages 1–8. IEEE, 2023.

[19] Zeming Lin, Halil Akin, Roshan Rao, Brian Hie, Zhongkai Zhu, Wenting Lu, Nikita Smetanin, Robert Verkuil, Ori Kabeli, Yaniv Shmueli, et al. Evolutionary-scale prediction of atomic-level protein structure with a language model. Science, 379(6637):1123–1130, 2023.

[20] Ahmed Elnaggar, Michael Heinzinger, Christian Dallago, Ghalia Rehawi, Yu Wang, Llion Jones, Tom Gibbs, Tamas Feher, Christoph Angerer, Martin Steinegger, et al. Prottrans: Toward understanding the language of life through self-supervised learning. IEEE transactions on pattern analysis and machine intelligence, 44(10):7112–7127, 2021.

[21] Ahmed Elnaggar, Hazem Essam, Wafaa Salah-Eldin, Walid Moustafa, Mohamed Elkerdawy, Charlotte Rochereau, and Burkhard Rost. Ankh: Optimized protein language model unlocks general-purpose modelling. arXiv preprint arXiv:2301.06568, 2023.

[22] Jack Hanson, Kuldip K Paliwal, Thomas Litfin, and Yaoqi Zhou. SPOT-Disorder2: improved protein intrinsic disorder prediction by ensembled deep learning. Genomics, Proteomics and Bioinformatics, 17(6):645–656, 2019.

[23] Claudio Mirabello and Björn Wallner. rawMSA: end-to-end deep learning using raw multiple sequence alignments. PloS one, 14(8):e0220182, 2019.

[24] Kui Wang, Gang Hu, Sushmita Basu, and Lukasz Kurgan. flDPnn2: Accurate and Fast Predictor of Intrinsic Disorder in Proteins. Journal of Molecular Biology, page 168605, 2024.

[25] Md Wasi Ul Kabir and Md Tamjidul Hoque. DisPredict3. 0: Prediction of intrinsically disordered regions/proteins using protein language model. Applied Mathematics and Computation, 472:128630, 2024.

[26] John Jumper, Richard Evans, Alexander Pritzel, Tim Green, Michael Figurnov, Olaf Ronneberger, Kathryn Tunyasuvunakool, Russ Bates, Augustin Žídek, Anna Potapenko, et al. Highly accurate protein structure prediction with alphafold. nature, 596(7873):583–589, 2021.

[27] Kiersten M Ruff and Rohit V Pappu. Alphafold and implications for intrinsically disordered proteins. Journal of molecular biology, 433(20):167208, 2021.

[28] Roshan Rao, Joshua Meier, Tom Sercu, Sergey Ovchinnikov, and Alexander Rives. Transformer protein language models are unsupervised structure learners. bioRxiv, 2020.

[29] Gang Hu, Akila Katuwawala, Kui Wang, Zhonghua Wu, Sina Ghadermarzi, Jianzhao Gao, and Lukasz Kurgan. flDPnn: Accurate intrinsic disorder prediction with putative propensities of disorder functions. Nature Communications, 12(1):4438, Jul 2021.

[30] András Hatos, Borbála Hajdu-Soltész, Alexander M Monzon, Nicolas Palopoli, Lucía Álvarez, Burcu Aykac-Fas, Claudio Bassot, Guillermo I Benítez, Martina Bevilacqua, Anastasia Chasapi, et al. DisProt: intrinsic protein disorder annotation in 2020. Nucleic acids research, 48(D1):D269–D276, 2020.

[31] Damiano Piovesan, Francesco Tabaro, Ivan Mic?etić, Marco Necci, Federica Quaglia, Christopher J Oldfield, Maria Cristina Aspromonte, Norman E Davey, Radoslav Davidović, Zsuzsanna Dosztányi, et al. DisProt 7.0: a major update of the database of disordered proteins. Nucleic acids research, 45(D1):D219–D227, 2017.

[32] Maria Cristina Aspromonte, Maria Victoria Nugnes, Federica Quaglia, Adel Bouharoua, Silvio CE Tosatto, and Damiano Piovesan. DisProt in 2024: improving function annotation of intrinsically disordered proteins. Nucleic Acids Research, 52(D1):D434–D441, 2024.

[33] Ying Huang, Beifang Niu, Ying Gao, Limin Fu, and Weizhong Li. CD-HIT Suite: a web server for clustering and comparing biological sequences. Bioinformatics, 26(5):680–682, 2010.

[34] Martin Steinegger and Johannes Söding. MMseqs2 enables sensitive protein sequence searching for the analysis of massive data sets. Nature biotechnology, 35(11):1026–1028, 2017.

[35] Felix Kallenborn, Alejandro Chacon, Christian Hundt, Hassan Sirelkhatim, Kieran Didi, Christian Dallago, Milot Mirdita, Bertil Schmidt, and Martin Steinegger. GPU-accelerated homology search with MMseqs2. bioRxiv, pages 2024–11, 2024.

[36] Chloé Dequeker, Elodie Laine, and Alessandra Carbone. INTerface Builder: A fast protein–protein interface reconstruction tool. Journal of Chemical Information and Modeling, 57(11):2613–2617, 2017.

[37] Simon J Hubbard and Janet M Thornton. Naccess. Computer Program, Department of Biochemistry and Molecular Biology, University College London, 2(1), 1993.

[38] Takuya Akiba, Shotaro Sano, Toshihiko Yanase, Takeru Ohta, and Masanori Koyama. Optuna: A next-generation hyperparameter optimization framework, 2019.

[39] I Loshchilov. Decoupled weight decay regularization. arXiv preprint arXiv:1711.05101, 2017.

[40] Zhenling Peng, Jing Yan, Xiao Fan, Marcin J Mizianty, Bin Xue, Kui Wang, Gang Hu, Vladimir N Uversky, and Lukasz Kurgan. Exceptionally abundant exceptions: comprehensive characterization of intrinsic disorder in all domains of life. Cellular and Molecular Life Sciences, 72:137–151, 2015.

